# Epilogos: information-theoretic navigation of multi-tissue functional genomic annotations

**DOI:** 10.1101/2025.06.18.660301

**Authors:** Jacob Quon, Alex Reynolds, Nalu Tripician, Eric Rynes, Athanasios Teodosiadis, Manolis Kellis, Wouter Meuleman

## Abstract

Functional genomics data, such as chromatin state maps, provide critical insights into biological processes, but are hard to navigate and interpret. We present Epilogos to address this challenge by offering a simple information-theoretic framework for large-scale visualization, navigation and interpretation of functional genomics annotations, and apply it to over 2,000 genome-wide chromatin state maps in human and mouse. We construct intuitive visualizations of multi-tissue chromatin state maps, prioritize salient genomic regions, identify group-wise differential regions, and enable rapid similarity search given a region of interest. To facilitate usability, we provide a purpose-built web-based browser interface (http://epilogos.net) alongside open-source software for community access and adoption.

## Main

High-throughput techniques for profiling epigenomic and transcriptional state across the human genome have generated rich genome-wide signal maps of histone modifications^1^, chromatin accessibility^2^, gene expression^3^, DNA methylation^4^, and chromatin regulator occupancy^5^. Each one of these maps captures distinct facets of genomic function, and thus multiple tracks are often integrated to describe the functional state of a genomic region within a given biological setting^5,6^. Numerous computational tools have been developed to translate these multiple epigenomic and functional genomics tracks into rich annotation labelings, including the chromatin states^7,8^ learned by tools such as ChromHMM^9^, SegWay^10^ and IDEAS^11^ (**Fig. 1a, Supp. Fig. 1a-c**).

**Figure 1 -.**
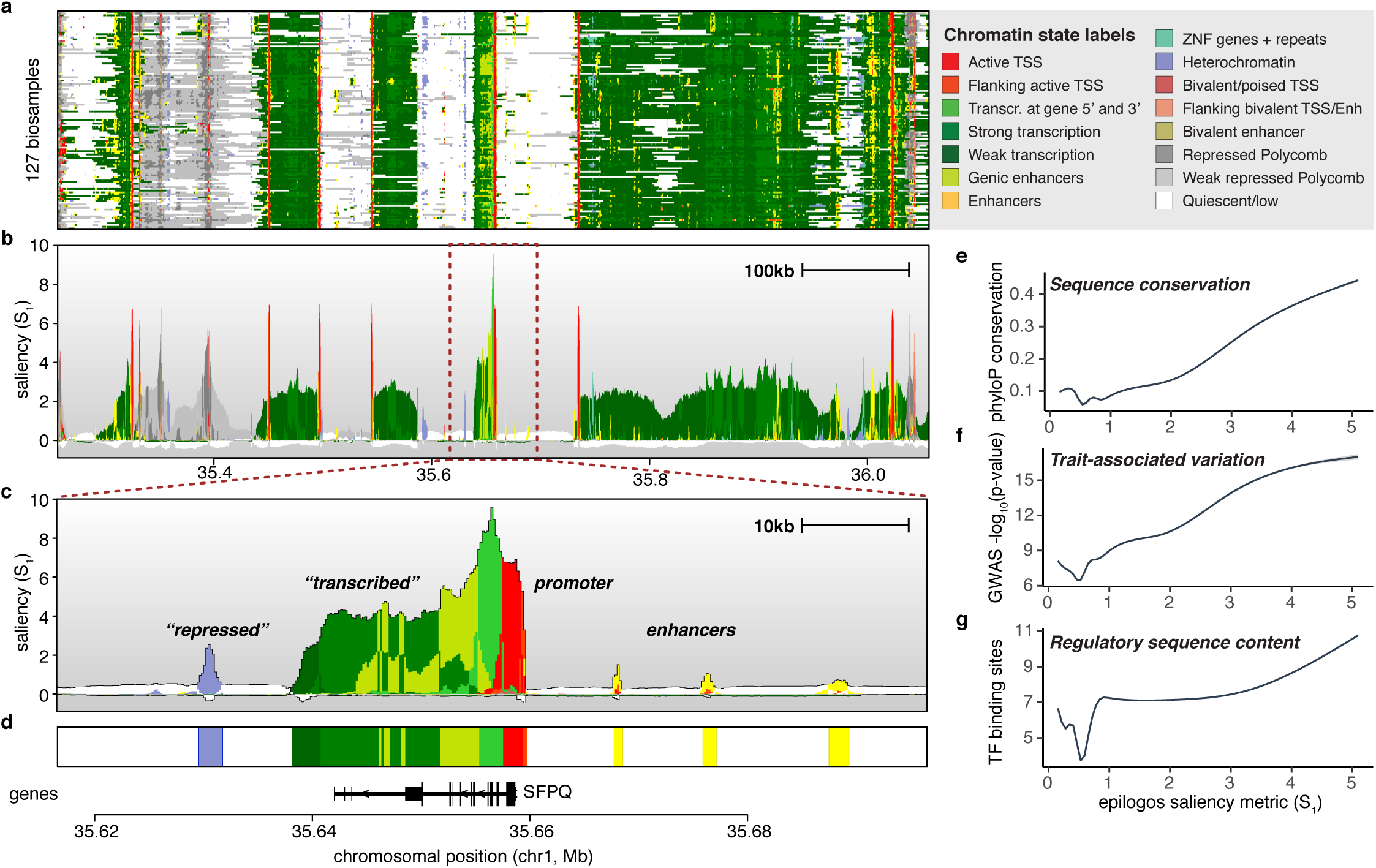
Summarization of chromatin state annotations using Epilogos. (**a**) Chromatin state annotations (15-state ChromHMM model) across 127 reference epigenome biosamples^43^ for an 800 kbp region on human chromosome 1. Colors indicate the various chromatin states as labeled on the right. (**b**) Epilogos scores for the same region and dataset as shown in (a), indicating levels of saliency, or relative entropy, for each genomic position. (**c**) Zoomed-in 80 kbp view around the SFPQ gene indicated by the dark red dashed rectangle in (b), highlighting a variety of regions highlighted by Epilogos, including those related to gene expression, regulation and repression. (**d**) Epilogos consensus track of chromatin states across all 127 biosamples, based on the maximal saliency state at each position. (**e**) Epilogos saliency scores correlate with the level of evolutionary sequence conservation, often used as an indicator for functional relevance. (**f**) Regions with high saliency scores capture genetic variants with stronger GWAS trait associations. (**g**) High saliency regions are enriched in regulatory signals encoded in the underlying genomic sequence.

Annotations derived from such tools have proven invaluable for identifying putative regulatory control regions in the human genome^12^, for interpreting genetic variation associated with human disease^13^, and for studying the dynamics of epigenomic information across cell types^14^ and development^15,16^. They have been widely adopted at two extremes in complexity, from visualization of individual regions, with one track representing each tissue or cell type^14,17^ (**Fig. 1a**), to calculations of aggregate statistics genome-wide, such as enrichment of disease-associated genetic variation in cell type-specific enhancers^13^. However, in between these two extremes, there is a notable lack of tools for studying such annotations across large numbers of biosamples or genomic regions. These tools are essential for discovering common and differential patterns and, more generally, for prioritizing and identifying regions of interest across biosamples. Such tools should quantitatively highlight salient patterns across datasets while remaining scalable enough to handle millions of regulatory elements across the genome and thousands of tissue and cell types.

### General Epilogos framework

To address these challenges, we developed Epilogos, a general framework for evaluating and utilizing the information content in functional genomics data. Epilogos builds upon information-theoretic tools traditionally applied to amino-acid or nucleotide sequences to model and detect common sequence patterns, or motifs^18^. Such motifs are generally summarized using sequence logos, which aggregate information across many individual sequences. We apply and extend this notion to genome annotation labelings (such as chromatin states), by considering an alphabet of such labels (instead of amino-acids or nucleotides) across multiple sequences representing biosamples (instead of molecules). In particular, we define saliency metrics to quantify the informativeness (non-randomness, unexpectedness) of a genomic region based on its labels across biosamples, relative to genome-wide annotation patterns (**Methods**).

Applying saliency measures genome-wide to chromatin state labels across biosamples results in saliency levels for each of the *n* labels at each nucleotide position. These levels quantify observed chromatin states relative to a background of expected (average) genome-wide occurrences (**Supp. Fig. 1d, Methods**). Analogous to sequence logos, chromatin state labels can be visualized in stacked representations, ordered and scaled by their relative saliency levels (**Fig. 1b**). Summed over all *n* labels, the total height of these stacks represents the overall per-position data saliency. The saliency levels provided by Epilogos readily highlight genomic regions with chromatin state configurations deviating from genome-wide expected patterns, for instance along gene bodies (**Fig. 1c**). Beyond such well-annotated regions, Epilogos highlights generally well-defined but poorly annotated regions such as repressive and enhancer elements (**Fig. 1c**).

This approach provides insights that are not immediately apparent from the hundreds of per-biosample chromatin state tracks (**Fig. 1a**), or from the hundreds to thousands of raw ChIP-seq datasets that constitute a chromatin state model. Beyond relative saliency levels for each of *n* labels at any genomic position, Epilogos facilitates the construction of a reference or consensus epigenome from a large number of individual epigenomes, by selecting the most salient state at each position (**Fig. 1d**). Note that this is different from choosing the dominant state across biosamples at each position (**Supp. Fig. 1e**). Indeed, the consensus highlights many more of the rare, often regulatory, chromatin states that are overlooked in averaged chromatin state occurrences (**Supp. Fig. 1f**) and per-position dominant states (**Supp. Fig. 1g-h**).

To validate the biological relevance of Epilogos saliency scores, we compared them with three independent sources of functional genomic information, namely (i) evolutionary sequence conservation, (ii) trait-associated genetic variation, and (iii) regulatory sequence content (**Methods**). First, Epilogos saliency scores strongly correlate with evolutionary sequence conservation across 100 vertebrate species (**Fig. 1e**), demonstrating its ability to prioritize conserved regions without explicit conservation data. Second, highly salient regions are enriched for genetic variants with strong trait associations (**Fig. 1f**), as quantified by the maximal trait association strength across traits included in the UK Biobank. Third, higher saliency scores identify regions with increased regulatory potential, evidenced by large numbers of distinct transcription factor binding motifs (**Fig. 1g**). Together, these results demonstrate the utility of Epilogos saliency scores for quantitatively prioritizing biologically relevant genomic regions in a data-driven manner.

### Differential analysis between groups of biosamples

In addition to representing a single group of biosamples, Epilogos enables direct pairwise comparisons between groups of biosamples. For example, grouping biosamples by donor sex—regardless of cell or tissue type—reveals significant epigenomic differences at loci central to X-inactivation^19^. These include the long non-coding RNA XIST (X-inactive specific transcript)^20^, as well as the X-linked tandem repeats FIRRE (Functional Intergenic Repeating RNA Element)^21^ and ICCE (Inactive-X CTCF-binding Contact Element)^22^ (**Fig. 2a**). With the increasing availability of genome-wide epigenomic datasets^23^, we extend this concept to unbiased, genome-wide comparisons between biosample groups (**Methods**). While sex-based differences are largely confined to distinct loci on chromosome X (**Fig. 2b-c, Supp. Fig. 2a**), other biosample groupings reveal differential loci distributed across the genome (**Fig. 2d, Supp. Fig. 2b-d**).

**Figure 2 -.**
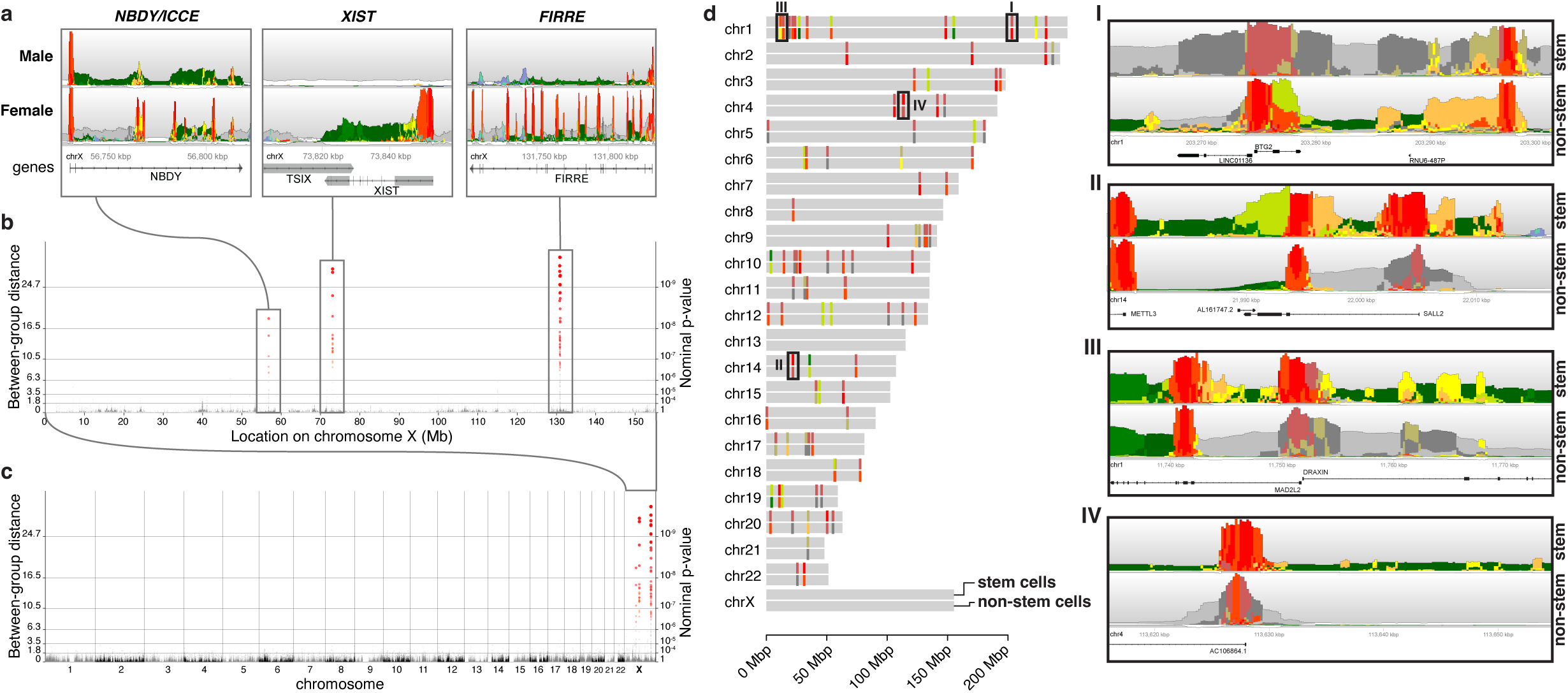
Pairwise comparison of biosample groups of interest. (**a**) Differential Epilogos profiles of biosamples obtained from male versus female donors, for several genic regions involved in the X-inactivation process (*NBDY/ICCE*, *XIST* and *FIRRE* genes). (**b**) Manhattan plot representation of differential Epilogos scores for male versus female donor biosamples on chromosome X, highlighting the three regions shown in (a). (**c**) Manhattan plot representation of differential male/female Epilogos scores for the entire human genome, illustrating that significant differences are confined to chromosome X only. (**d**) Genomic locations of the top 100 differentially scoring regions between stem cell biosamples and non-stem cell biosamples, with colors indicating highest scoring chromatin states in each. The following four regions are highlighted. (**I**) Anti-proliferation gene *BTG2*, inactive in stem cells (ranked #2/100). (**II**) Embryonic neurogenesis transcription factor *SALL2* (ranked #30/100). (**III**) Nervous system development gene *DRAXIN* (ranked #42/100). (**IV**) Due to its annotation-agnostic nature, Epilogos is able to identify differential but otherwise poorly characterized regions.

Among the top 100 differentially marked genomic regions between embryonic stem cells (ESCs) and non-stem cell biosamples (**Fig. 2d**), we identify loci associated with key regulators of cell cycle and development. These include BTG2 (#2/100, inactive in ESCs), an anti-proliferation gene^24^ (**Fig. 2d, panel I**), and SALL2 (#30, active in ESCs), a transcription factor essential for embryonic neurogenesis^25,26^ (**Fig. 2d, panel II**). Additional regions include MAD2L2 and DRAXIN (#42, active in ESCs), which play roles in germ cell and nervous system development^27,28^ (**Fig. 2d, panel III**). Notably, because Epilogos operates independently of gene annotations, it identifies differential epigenomic profiles in both coding and non-coding regions, such as an antisense long noncoding RNA (lncRNA) implicated in gastrulation^29^ (**Fig. 2d, panel IV**). Similarly, differential analysis between cancer and non-cancer biosamples (**Supp. Fig. 2b**) highlights key oncogenic and tumor-suppressor loci. Notable examples include BRCA1 (#3), KIF4A (#4), MYC (#29), and ZFAS1 (#1), a long non-coding RNA linked to tumor progression and metastasis^30^. Differentially marked loci are also enriched for ribosomal protein genes^31,32^ and small nucleolar RNA host genes (SNHGs)^33^, both of which are associated with cancer metastasis and poor prognosis. Given the vast amount of publicly available chromatin state data, Epilogos can be used to study epigenomic patterns specific to any group of biosamples of interest, such as immune cells (**Supp. Fig. 2c**) or neural cell types (**Supp. Fig. 2d**).

In contrast to existing methods^34–37^, Epilogos explicitly accounts for genome-wide chromatin state frequencies, providing an information-theoretic measure of state and region saliency. Unlike previous approaches, it is not restricted to pre-defined genomic loci^34^, chromatin states^36^, or biosamples^35^, ensuring a fully data-driven and unbiased approach to differential analysis. Identified differential regions are paired with an intuitive visual representation, combining both the magnitude of epigenomic differences and the functional significance of the involved chromatin states. This approach not only provides a powerful tool for prioritizing differential regions but also offers a framework that can integrate and refine existing^38^ or future methods for quantifying pairwise differences.

### Similarity search across the genome

Navigating the functional organization of the human genome is challenging due to its immense complexity. However, large-scale visualizations of chromatin state data, such as those generated by Epilogos, reveal remarkably stereotypical genome-wide patterns. For instance, genes are typically marked by promoter-related states (e.g. ‘TssA’), followed by transcribed states if the gene is expressed (e.g. ‘Tx’), and clusters of regulatory elements such as enhancers (e.g. ‘Enh’) (**Fig. 1**). These recurring chromatin signatures serve as visual anchors, facilitating navigation of the epigenome.

To facilitate genome exploration, we developed a simple similarity-based search in Epilogos to identify regions with epigenomic patterns similar to a given query region. Using a combination of information-preserving search space reduction and similarity-based scoring (**Methods**), the approach allows for the identification of regions that are numerically and visually similar to a given genomic location of interest.

To illustrate this approach, we analyzed a 250kb genomic region on human chromosome 10 (**Fig. 3a**), identifying multiple highly similar regions across the genome (**Fig. 3b-d**). For instance, a 10kb region encompassing the promoter of the CHST3 gene has several matching regions on different chromosomes (**Fig. 3b**). Similarly, the largely bivalent promoter of the SPOCK2 gene has highly similar matches to other bivalent promoters across multiple chromosomes (**Fig. 3d**). In the same larger region, a 25kb enhancer-dense region has close matches with other enhancer-rich loci elsewhere in the genome (**Fig. 3c**). Indeed, this search approach is scalable to regions of arbitrary size, ranging from 5kb to 100kb (**Supp. Fig. 3**). Taken together, this approach enables rapid identification of regions with stereotypical chromatin patterns at a variety of scales, providing a simple, intuitive, and scalable way to explore the human epigenome.

**Figure 3 -.**
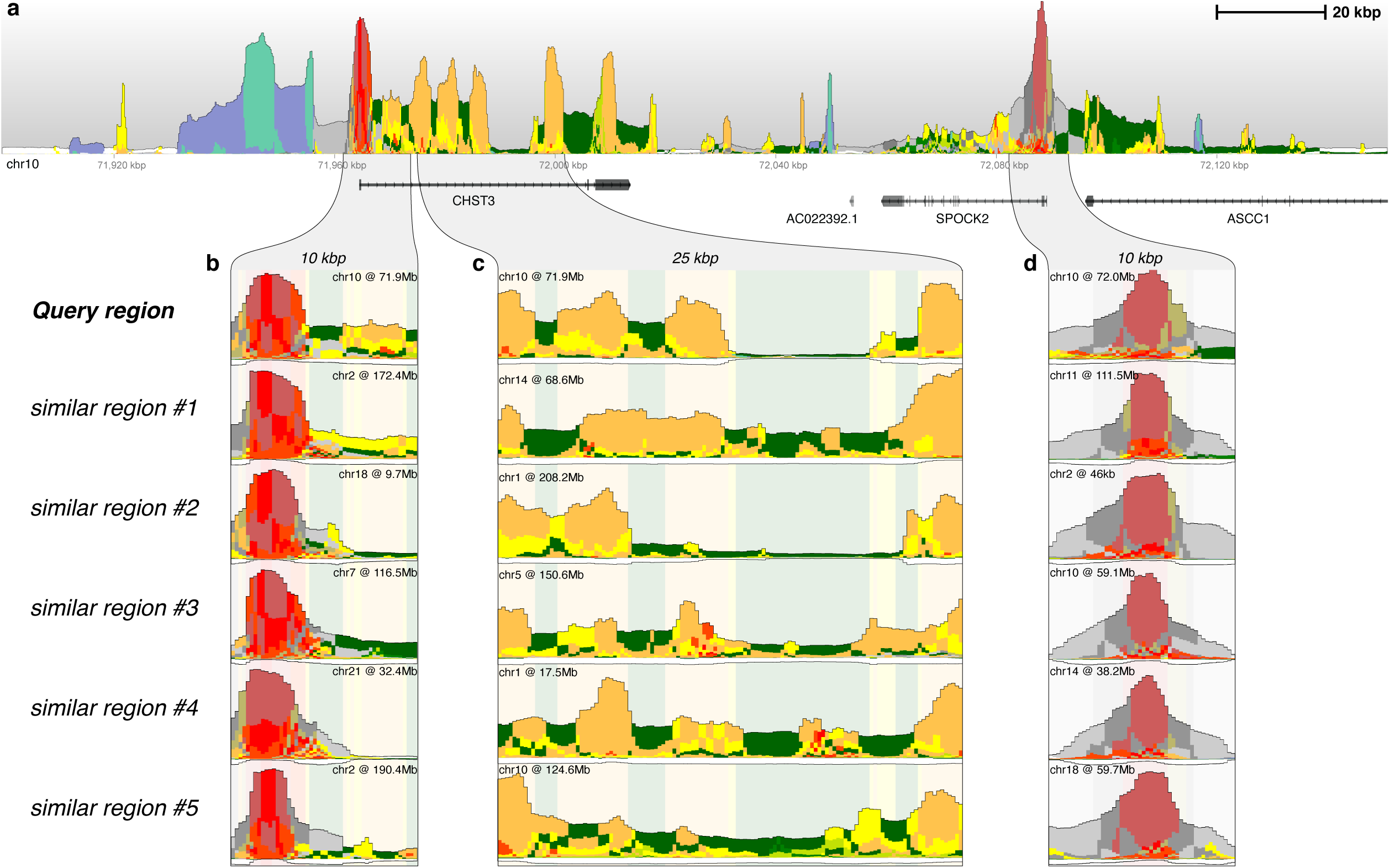
Similarity search based on Epilogos saliency scores. (**a**) A 250 kbp region on human chromosome 4 showing a wide variety of chromatin states and saliency score levels. (**b**) A 10 kbp query region found in (a), along with its top 5 most similar regions across the entire genome, showing similar promoter architectures. (**c**) A 25 kbp query region and its top 5 most similar regions, highlighting a repeated pattern of enhancer clusters across the genome. (**d**) A 10 kbp query region and its top 5 most similar regions, illustrating the recurrence of a strong bivalent promoter and its Polycomb repressed flanks. Background colors in (b-d) are based on the consensus Epilogos calls of the query regions.

### Epilogos data browser and resources

Here, we applied the Epilogos framework to four large reference datasets, encompassing over 2,000 genome-wide chromatin state maps from human^12,39^ and mouse^40^ (**Table 1**). The largest of these is a set of 1,698 human epigenomes collected, curated and processed by the International Human Epigenome Consortium (IHEC). To visualize these data, we developed an interactive browser built on top of the HiGlass^41^ platform, originally designed for visualizing large Chromosome Conformation Capture (3C) experiment data. In collaboration with the HiGlass developers, we extended its capabilities to support chromatin state annotations and Epilogos scores. The result is a fast, lightweight, and fully web-based browser (https://epilogos.net), that supports a wide range of interactive features (**Fig. 4a**).

**Figure 4 -.**
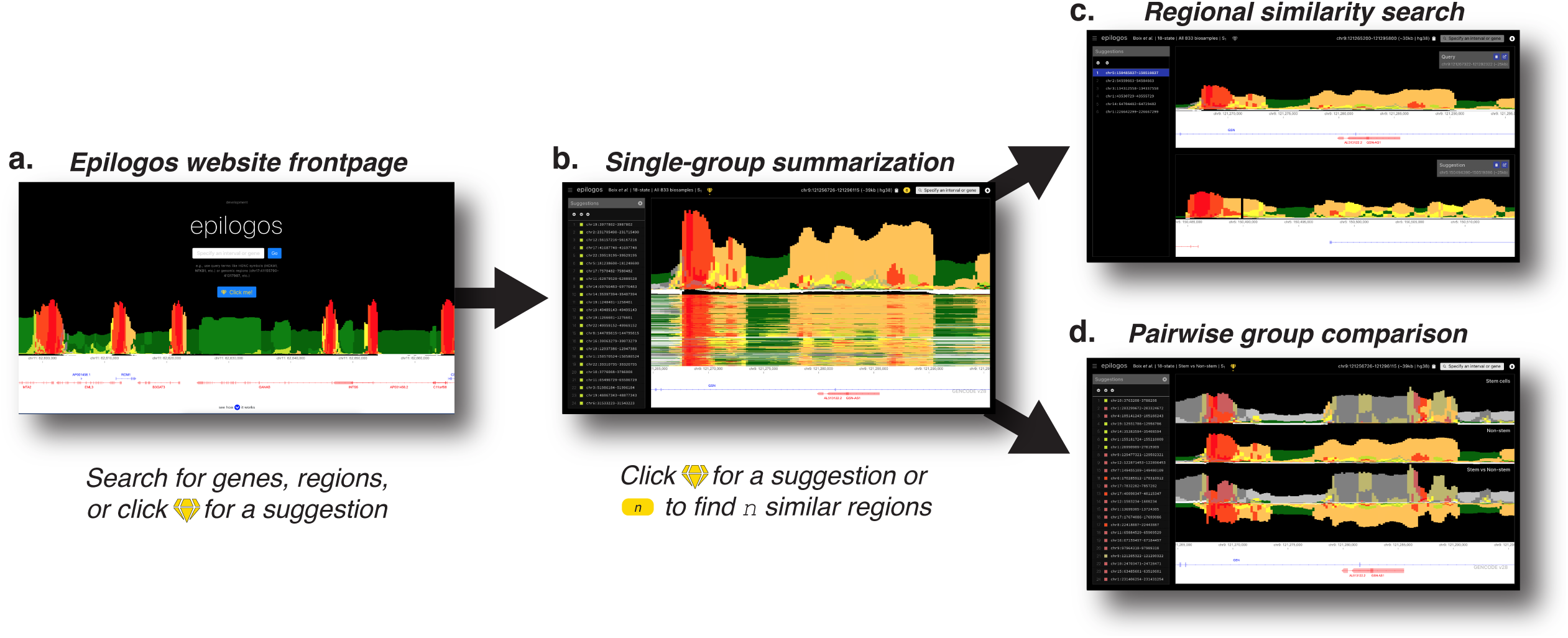
Epilogos website functionality and flow. (**a**) Epilogos website frontpage, with functionality to search for genes and regions of interest, or receive a random high-saliency suggestion. (**b**) Main Epilogos view, summarizing chromatin state tracks across biosamples into a single Epilogos track. (**c**) Similarity search allows for the retrieval of visually similar regions based on (b). (**d**) Pairwise comparison view, highlighting differences between biosample groups of interest.

**Table 1 -.**
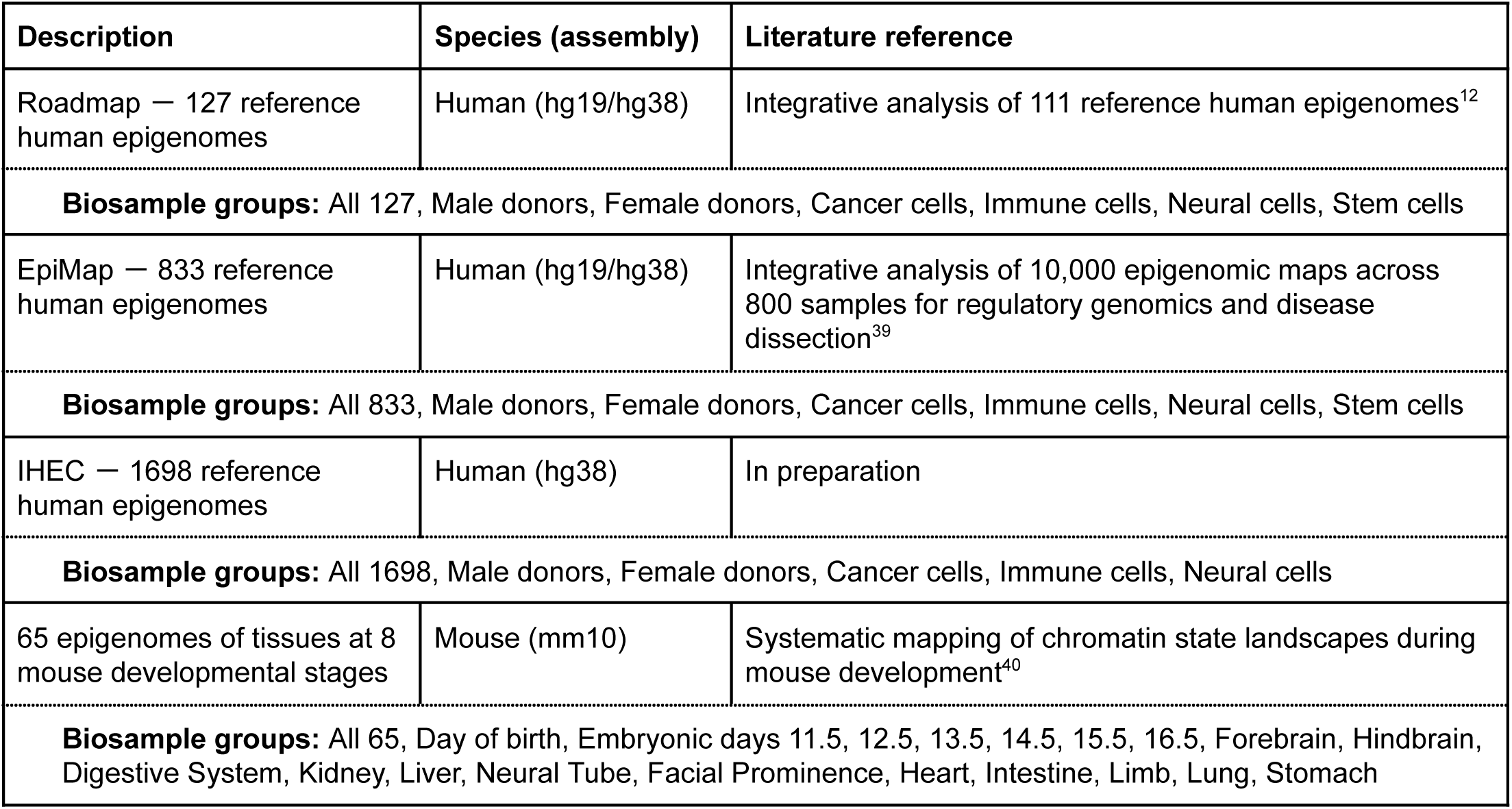
Available datasets.

| Description | Species (assembly) | Literature reference |
| --- | --- | --- |
| Roadmap — 127 reference human epigenomes | Human (hg19/hg38) | Integrative analysis of 111 reference human epigenomes<sup>12</sup> |
| **Biosample groups:** All 127, Male donors, Female donors, Cancer cells, Immune cells, Neural cells, Stem cells |  |  |
| EpiMap — 833 reference human epigenomes | Human (hg19/hg38) | Integrative analysis of 10,000 epigenomic maps across 800 samples for regulatory genomics and disease dissection<sup>39</sup> |
| **Biosample groups:** All 833, Male donors, Female donors, Cancer cells, Immune cells, Neural cells, Stem cells |  |  |
| IHEC — 1698 reference human epigenomes | Human (hg38) | In preparation |
| **Biosample groups:** All 1698, Male donors, Female donors, Cancer cells, Immune cells, Neural cells |  |  |
| 65 epigenomes of tissues at 8 mouse developmental stages | Mouse (mm10) | Systematic mapping of chromatin state landscapes during mouse development<sup>40</sup> |
| **Biosample groups:** All 65, Day of birth, Embryonic days 11.5, 12.5, 13.5, 14.5, 15.5, 16.5, Forebrain, Hindbrain, Digestive System, Kidney, Liver, Neural Tube, Facial Prominence, Heart, Intestine, Limb, Lung, Stomach |  |  |

We created Epilogos tracks for each full reference dataset, as well as for many biosample subsets sharing common characteristics (**Fig. 4b**). For example, these include Epilogos tracks restricted to immune cell types or to biosamples only from male donors (**Table 1**). We used such groupings to perform pairwise comparisons between biosample sets of interest, such as immune vs. non-immune cells or male vs. female donors (**Fig. 4c**). The similarity-based search described earlier is integrated into the browser and supports queries at multiple genomic scales, from 5 kb to 100 kb (**Fig. 4d**). Throughout the browsing experience, users receive suggestions for regions to explore next, based on either high Epilogos saliency scores or pattern similarity to their region of interest (**Fig. 4**).

In addition to interactive browsing, the Epilogos Browser allows users to download data for selected genomic regions and export publication-quality figures. We provide the browser as a public resource, along with open-source implementations of the Epilogos browser (https://github.com/meuleman/epilogos-web), the core Epilogos framework (https://github.com/meuleman/epilogos), and the `epilogos` Python package, available via the Python Package Index (PyPI).

## Discussion

Segmentation of genome-wide functional data has become a cornerstone of large-scale genomic analysis, enabling the annotation of chromatin states and regulatory elements across diverse biosamples. While methods for generating these annotations have steadily improved, their interpretation - especially across thousands of epigenomes - remains challenging and often requires expert knowledge. With large consortium efforts like ENCODE and IHEC producing thousands of epigenomes, scalable and accessible tools for data analysis are increasingly essential.

Epilogos addresses this challenge by reducing the dimensionality of large genome-wide annotations while preserving key biological signals. It enables rapid visualization, intuitive summarization, and pairwise comparisons across biosample groups. Without relying on any external genomic annotation, Epilogos reveals both well-characterized and previously underappreciated genomic regions, including many non-coding and potentially novel regulatory loci. It also supports similarity-based navigation, leveraging the stereotypical nature of chromatin state patterns to enable efficient exploration of related regions.

Epilogos builds on well-established principles from the field of Information Theory to compute saliency scores that quantify the informativeness of chromatin state patterns at each genomic position. While this work focuses on chromatin state segmentations, the framework is broadly applicable to any genome-wide annotation of countable events, including gene annotations (e.g. exons, introns, non-coding RNAs), evolutionary conservation (e.g. mammalian-conserved), and transcription (e.g. cell-type-specific expression patterns).

A previously proposed “functionality score”^42^ appears superficially similar to Epilogos in its visual representation. However, it relies on evolutionary sequence conservation to scale chromatin state stacks, implicitly assuming that only conserved regions are functionally relevant. In contrast, Epilogos derives saliency scores solely from the epigenomic data itself, offering a more data-driven and unbiased approach. Importantly, the functionality score yields quantitatively identical results across different biosample groups, whereas Epilogos allows group-specific interpretation by accounting for the information content of each. Moreover, the functionality score provides limited resolution into contribution of individual chromatin states, whereas Epilogos quantifies how unexpected or surprising observed chromatin configurations are relative to genome-wide expectations. Although evolutionary conservation could be layered on top of Epilogos as a secondary score, we find that Epilogos saliency already tracks well with conservation (**Fig. 1e**).

Taken together, Epilogos adds an essential interpretive layer to large-scale functional genomics pipelines. It enables researchers to explore, summarize, and compare massive datasets in a straightforward and interpretable manner. Just as sequence logos represent the motifs of primary DNA sequences, Epilogos offers an analogous representation for chromatin state and other functional genomics-based annotations.

We demonstrate that Epilogos readily identifies salient regions of interest and supports comparative analysis across biosample groups. Its intuitive visual output and flexible framework provide a scalable and principled tool for genome interpretation. These ideas contribute to an emerging paradigm we term augmented genomics: a strategy that combines large-scale visualization with data-driven inference to support genome scientists in navigating the rapidly growing landscape of functional genomics data.

## Acknowledgements

Soheil Feizi and Angela Yen for helpful discussions. Apostolos Papadopoulos for an early prototype of the web-interface. Research reported in this publication was supported by the National Human Genome Research Institute of the National Institutes of Health under Award Number R35HG011317.

## Author information

W.M. conceived of the project. W.M., J.Q. and N.T. contributed to method development. J.Q., E.R. and W.M. implemented the software. A.R. developed the web-based browser. J.Q., A.R., E.R., N.T., A.T., M.K. and W.M. analyzed the data. W.M., M.K. and J.Q. wrote the manuscript with input from all coauthors.

## Supplementary Materials

Supplementary Figures are available in the online version of the paper.

## Methods

### Chromatin state annotations and data representation

We use genome-wide chromatin state annotations that assign a categorical label to each 200-bp genomic segment. These labels represent chromatin states^9,10^ and summarize combinations of epigenomic features - including DNA accessibility, histone modifications, and DNA methylation - into a single annotation track per biosample (**Fig. 1a**). Chromatin states define a shared vocabulary of recurrent epigenomic mark combinations across cell types and yield an interpretable segmentation of the genome into discrete intervals.

We use chromatin state annotations generated by ChromHMM^9^, modeling state assignments based on the co-occurrence of histone marks (e.g., H3K27me3, H3K4me1, etc). A key parameter to ChromHMM is the number of discrete states to learn; different models yield different sets of states that correspond to features such as promoters, enhancers, transcribed regions, and repressed or quiescent domains. In this work, we include models with 15, 18 and 25 states. Although we use ChromHMM’s nucleosome-sized 200-bp resolution, Epilogos is compatible with segmentations at any resolution.

Collections of chromatin state tracks across many biosamples serve as input to Epilogos. As part of this work, we generated and analyzed Epilogos data from the following datasets:

- Roadmap Epigenomics Project^43^: 15-, 18-, and 25-state annotations across 127 human biosamples (https://compbio.mit.edu/roadmap/)
- EpiMap project^23^: 18-state annotations for 833 human biosamples (https://compbio.mit.edu/epimap/)
- International Human Epigenome Consortium (IHEC): 18-state annotations for 1,698 human biosamples (https://epigenomesportal.ca/ihec)
- Mouse epigenomes^44^: 15-state annotations for 65 mouse biosamples (https://www.encodeproject.org/)

### Information-theoretic metrics of data saliency

Information theory provides a natural framework for quantifying saliency in labeled functional genomics data. Analogous to the use of sequence logos to describe aligned DNA or protein sequences, we apply similar principles to genome annotation tracks composed of categorical labels (e.g., chromatin states). To address the general challenge of region prioritization, we define three saliency metrics - S1, S2, and S3 - that measure the informativeness (i.e., non-randomness or unexpectedness) of a genomic region based on observed annotation patterns across biosamples.

#### Relative entropy (S_1_)

As a baseline, we compute saliency using *relative entropy*, also known as Kullback–Leibler (KL) divergence. For each chromatin state label i, the saliency score is defined as:

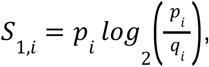

where *p_i_* is the observed frequency of label *i* at a genomic position (across biosamples), and *q_i_* is its expected frequency genome-wide. This score quantifies the information gain from observing label *i* relative to genome-wide expectation. This metric naturally downweights common, less informative states (e.g. quiescent or transcribed chromatin states, which collectively cover more than 80% of the genome) and upweights more rare labels (e.g. enhancer states, promoter states, which comprise less than 5%) (**Supp. Fig. 1d**). We refer to this measure as S_1_ and use it as the default metric throughout this work due to its simplicity, interpretability, and computational efficiency.

#### State co-occurrence-adjusted entropy (S_2_)

The S_2_ metric builds on S_1_ by incorporating pairwise co-occurrence patterns between chromatin states. This accounts for differences in the sample-specificity of states. For instance, promoter states tend to be consistently observed across biosamples, whereas enhancer states are more cell-type-specific. Observing multiple enhancer states across biosamples at the same genomic position is thus more surprising than observing multiple promoter states.

To capture this, we compute a pairwise co-occurrence matrix across all biosamples. For each label i, the saliency score is calculated as:

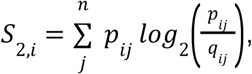

where *p_ij_* is the observed co-occurrence frequency of labels *i* and *j*, and *q_ij_* is the expected co-occurrence under a genome-wide background model. This formulation allows S_2_ to highlight regions where combinations of states (e.g. enhancer states in multiple biosamples) occur more frequently than expected by chance.

#### State and biosample co-occurrence-adjusted entropy (S_3_)

The S_3_ metric extends S_2_ by incorporating biosample-specific relationships. It models the likelihood of co-occurring chromatin state pairs across pairs of biosamples, reflecting that more similar biosamples are more likely to share chromatin state configurations.

The saliency score for each label *i* is defined as:

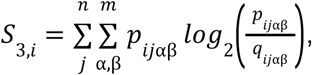

where *p_ijαβ_* is the frequency of observing label *i* in biosample *α* and label *j* in biosample *β* at a given position, and *q_ijαβ_* is the expected co-occurrence under a background model. This formulation captures more complex dependencies between chromatin state configurations and biosample relationships, at the cost of greater computational complexity.

#### Summary of saliency metrics

The three saliency metrics represent a progression from simple to more complex background models:

- S1: relative entropy of label frequency vs. genome-wide expectation
- S2: relative entropy adjusted for pairwise chromatin state co-occurrence
- S3: relative entropy adjusted for both state and biosample pairwise co-occurrence

All three metrics compute per-position and per-label saliency scores using label frequencies across the full set of biosample annotations. Unless otherwise specified, we apply S1 throughout this study for its balance of computational efficiency and biological interpretability.

### Epilogos saliency scores vs. orthogonal annotations

To assess the biological relevance of Epilogos saliency scores, we compared them against three independent sources of functional genomic information: evolutionary sequence conservation, trait-associated genetic variation, and regulatory sequence content (**Fig. 1e-g**). We first binned the genome into 100 equally sized groups based on Epilogos saliency scores computed across 127 human biosamples from the Roadmap Epigenomics Project. These “saliency bins” reflect increasing levels of per-position Epilogos saliency. For each bin, we computed the median saliency score and plotted it against each of the three orthogonal annotations as described below.

#### Sequence conservation

To evaluate sequence conservation, we obtained phyloP scores from the UCSC Table Browser (phyloP100way track), which represent nucleotide-level evolutionary conservation across 100 vertebrate species. We averaged phyloP scores across all positions in each saliency bin to obtain a conservation score per bin.

#### Trait-associated genetic variation

To quantify the relevance of saliency scores to common disease or complex trait variation, we used genome-wide association study (GWAS) summary statistics from the UK Biobank (2018 release). For each 200-bp genomic bin (matching the Epilogos resolution), we computed the maximum −log10(p-value) across all tested traits and variants. These values were then averaged across each saliency bin to derive a per-bin trait-association score.

#### Regulatory sequence content

To estimate regulatory sequence potential, we assessed transcription factor (TF) motif density. TF binding motifs were collected from the TRANSFAC, UNIPROBE, Taipale, and JASPAR databases, scanned across the human genome using FIMO (Find Individual Motif Occurrences) with a significance threshold of 10^-5^, and clustered based on similarity (http://www.mauranolab.org/CATO/weblogos/main.html). For each 200-bp bin, we counted the number of distinct motif clusters with at least one match. Finally, we averaged these motif cluster counts within each saliency bin to obtain a regulatory content score.

### Pairwise biosample group comparison

To identify regional differences in chromatin state configurations between two biosample groups, we compute a simple Signed Squared Euclidean Distance (SSED) metric between their Epilogos saliency profiles at each genomic position. The SSED is defined as the sum of squared differences between the saliency scores of chromatin states in two groups *A* and *B*:

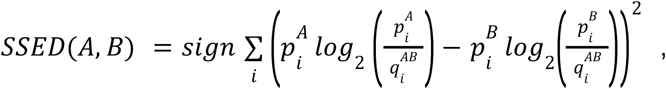

Where the sign is determined by which group has the higher total saliency (i.e., sum over all states). This formulation emphasizes positions where the distributions of chromatin states differ substantially between the groups, while preserving the direction of difference. Saliency scores are generated in the same manner as in the single group case (see S1 metric) with the exception that the expected frequencies are calculated jointly across both biosample groups to ensure a consistent background.

To assess the statistical significance of observed differences, we use a permutation-based approach. We repeatedly permute biosample group labels and recompute SSED scores and fit a generalized normal distribution. To avoid inflation of zero-valued bins, we exclude genomic bins in which all biosamples are assigned the quiescent state. For computational efficiency, we randomly repeatedly sample sets of 100,000 genomic bins across permutations and fit the distribution to each sampled set. The final null model is selected as the median fit across these samples, based on the negative log-likelihood. Empirical p-values are derived from this null model, and we apply the Benjamini-Hochberg procedure to control the False Discovery Rate (FDR) across genomic positions.

### Region recommendation and similarity search

Epilogos saliency scores provide a foundation for identifying and recommending functionally informative genomic regions. We implement two simple strategies to support this: (1) region recommendation based on global saliency, and (2) region-based similarity search to identify regions with comparable chromatin state saliency patterns.

#### Region recommendation

To identify generally salient genomic regions, we curate a non-overlapping set of high-saliency intervals using a greedy algorithm. Starting from individual high-scoring 200-bp bins (based on Epilogos saliency scores), the algorithm extends each bin asymmetrically to form a region of fixed size. The extension direction is chosen to maximize the aggregate saliency within the region, and overlapping regions are excluded to ensure independence. This results in a ranked list of salient genomic intervals. For each dataset presented in the Epilogos browser, we select the top 100 such regions and offer them as general recommendations. These regions can be browsed interactively via the “gem” icon in the browser interface. By default, recommended regions are 10 kb in size for single-group saliency visualizations, and 25 kb for pairwise comparisons.

#### Similarity search

In addition to static recommendations, Epilogos supports interactive similarity search based on a user-specified query region. This functionality is designed to accommodate the visual intuitions of users, who may perceive two regions as similar even if they contain small differences in alignment or spacing that would obscure exact one-to-one comparisons. Such differences do not necessarily influence the functional interpretation of data in a given genomic region. For instance, a slight difference in the spacing between two enhancer regions, or the precise length of a transcribed region flanked by regulatory regions. This human tolerance increases as larger genomic regions are being considered.

To allow for such flexible matching, we implement a scale-normalized data reduction approach. Each region of interest is first divided into 25 non-overlapping “blocks”, regardless of its original size. This is achieved via a max-pooling–like operation over consecutive 200-bp bins, ensuring that resolution scales with region length. The resulting vector of 25 block-level saliency scores serves as a compact, standardized representation of the region. We then compare this representation to those of all other regions genome-wide using a Euclidean distance metric. To identify highly similar regions, we apply a similarity threshold equal to half the mode of the genome-wide distance distribution, ensuring that only regions more similar than dissimilar to the query are returned. This simple yet robust approach enables users to discover regions with similar epigenomic configurations, accounting for approximate alignment and variation in chromatin state patterns, especially over larger genomic intervals.

## Figure Legends

**Supp. Figure 1 - Chromatin state models and saliency scores**

(**a-c**) A 15-state ChromHMM model as used across 127 reference epigenomes^12^, based on H3K4me1, H3K4me3, H3K27me3, H3K36me3 & H3K9me3 histone tail modification ChIP-seq data. Shown are (**a**) chromatin state labels and abbreviations, (**b**) emission matrix containing the combinatorial patterns of epigenomic marks per chromatin state and (**c**) enrichments of various genomic annotations and functional activity levels per chromatin state. (**d**) Average genome-wide chromatin state frequencies across all 127 epigenomes, used as background (expected) frequencies for calculating Epilogos saliency scores. (**e**) Consensus state frequencies, as determined by per-position maximum saliency states, shown alongside state frequencies as determined by choosing the dominant state at each position. (**f-g**) Comparison of consensus versus (f) average and (g) dominant state frequencies, highlighting the over-representation of regulatory and other non-quiescent states in the consensus states. (**h**) Comparison of per-position consensus versus dominant state frequencies, highlighting an over-representation of quiescent state calls in the dominant state frequencies.

**Supp. Figure 2 - Top 100 differential regions for various groupings**

**(a)** Statistically significant differential regions between male and female donor biosamples, highlighting only X-inactivation related genes and regions on chromosome X. (**b**) Differential regions between cancer and non-cancer biosamples, highlighting cancer related genes *BRCA1*, *ZFAS1*, *KIF4A*, & *MYC*. (**c**) Differential regions between immune and non-immune biosamples. (**d**) Differential regions between neural and non-neural biosamples.

**Supp. Figure 3 - Similarity search results across genomic scales**

**(a)** a 5kb promoter-like region on human chromosome 2 and its 5 most similar regions elsewhere in the genome. (**b**) a 10kb region with strong regulatory signals on chromosome 6 and its top 5 similar regions. (**c,d**) two separate 10kb regions on chromosomes 3 and 19, highlighting similarity across loosely organized clusters of enhancers (**c**) and regions overlapping zinc-finger genes (**d**, gene names indicated). (**e**) A large 100kb region in the HOXA locus, with functionally highly related matches in other hox loci, namely HOXD, HOXB and HOXC.

