## Supplementary figures and images for "Epilogos: information-theoretic navigation of multi-tissue functional genomic annotations"

### Supp Figure 1

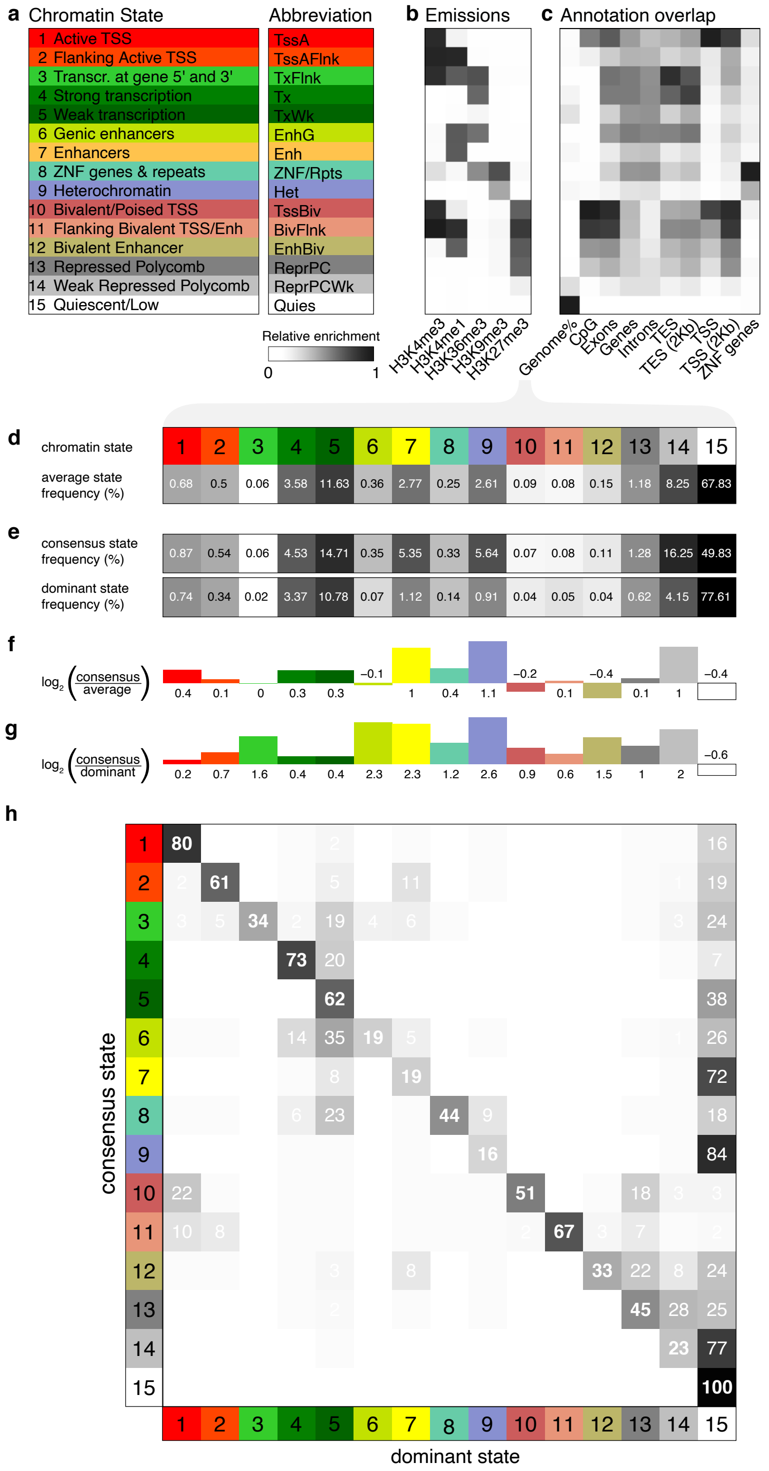

### Supp Figure 2

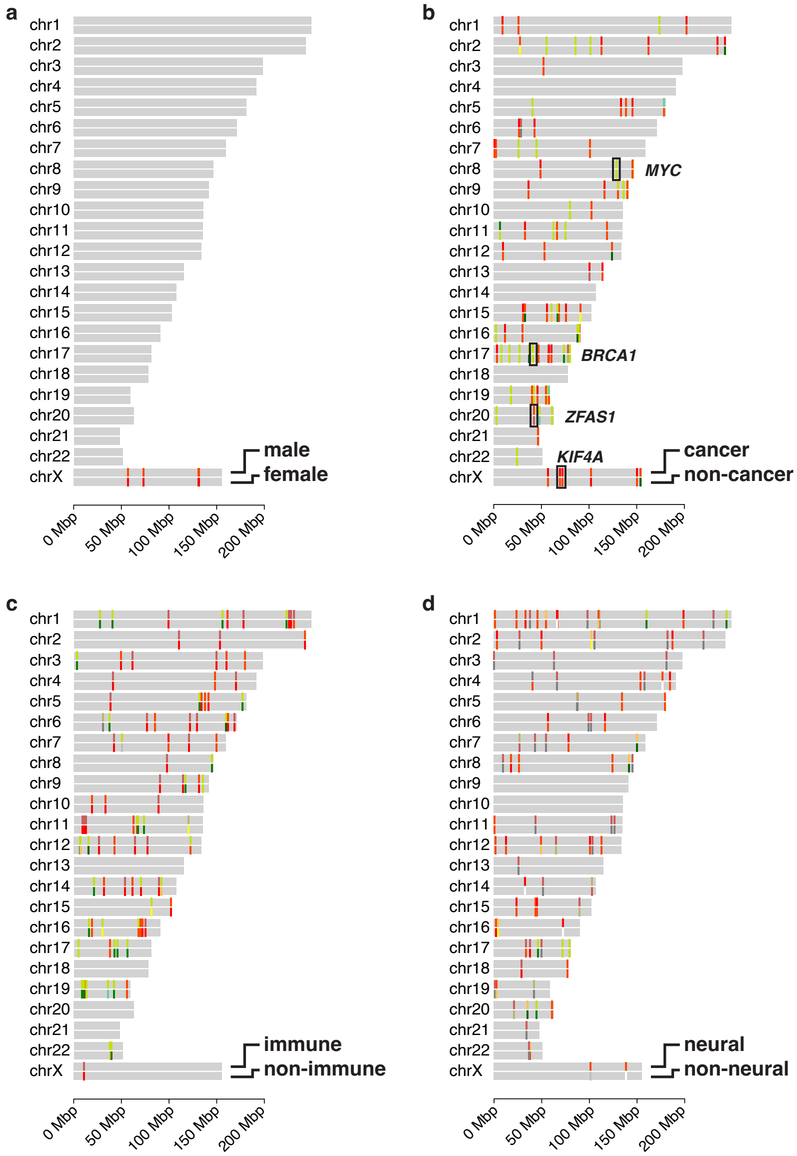

### Supp Figure 3

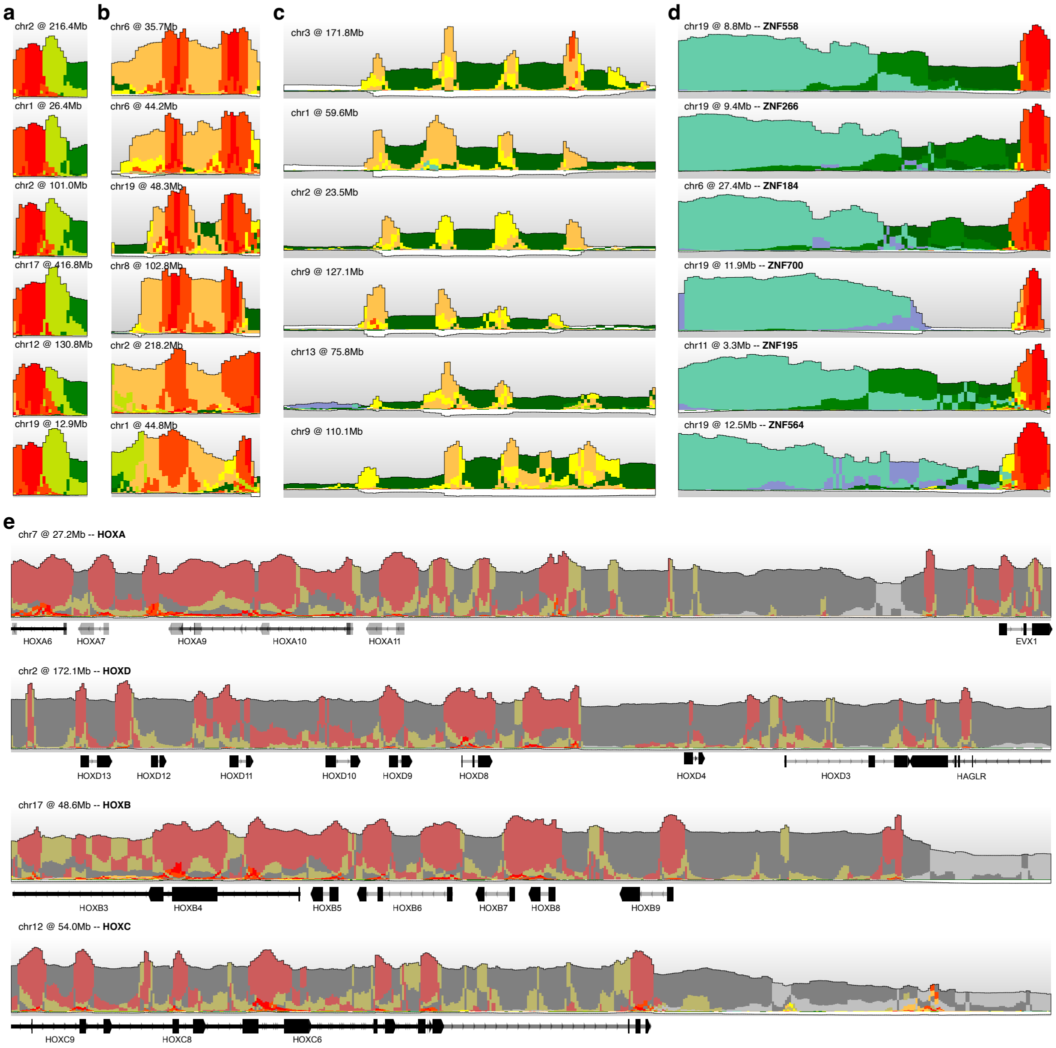
